# The kinetoplastid-infecting Bodo saltans virus (BsV), a window into the most abundant giant viruses in the sea

**DOI:** 10.1101/214536

**Authors:** Christoph M. Deeg, Cheryl-Emiliane T. Chow, Curtis A. Suttle

## Abstract

Giant viruses are ecologically important players in aquatic ecosystems that have challenged concepts of what constitutes a virus. Herein, we present the giant Bodo saltans virus (BsV), the first characterized representative of the most abundant giant viruses in the oceans, and the first klosneuvirus isolate, a subgroup of the *Mimiviridae* proposed from metagenomic data. BsV infects an ecologically important microzooplankton, the kinetoplastid *Bodo saltans*. Its 1.39 Mb genome is the largest described for the *Mimiviridae* and encodes 1227 predicted ORFs, including pathways for host-independent replication. Yet, much of its translational machinery has been lost, including all tRNAs. Essential genes are invaded by homing endonuclease-encoding self-splicing introns that may defend against competing viruses. Putative anti-host factors show extensive gene duplication via a genomic accordion indicating an ongoing evolutionary arms race and highlighting the rapid evolution and genomic plasticity that has led to genome gigantism and the enigma that is giant viruses.

## Introduction

Viruses are the most abundant biological entities on the planet and there are typically millions of virus particles in each ml of marine or fresh waters that are estimated to kill about 20% of the living material, by weight, each day in the oceans (Curtis A. Suttle, 2007). This has major consequences for global nutrient and carbon cycles, as well as for controlling the composition the planktonic communities that are the base of aquatic foodwebs. Although the vast majority of these viruses are less than 100 nm in diameter and primarily infect prokaryotes, it became increasingly clear that a subset of the viruses in aquatic ecosystems are comparative Leviathans that have been anecdotally classified as giant viruses.

The first giant virus that was isolated and described was for a marine microzooplankton that was misidentified as *Bodo* sp. (Garza, 1995), but which actually infected a protozoan from a different evolutionary branch of eukaryotes, *Cafeteria roenbergensis (Fischer, Allen, Wilson, & Suttle, 2010)*. However, it was the isolation and sequencing of mimivirus, a giant virus infecting *Acanthamoeba polyphaga* (La Scola, 2003; Raoult et al., 2004) that transformed our appreciation of the biological and evolutionary novelty of giant viruses. This led to an explosion in the isolation of different groups of giant viruses infecting *Acanthamoeba* spp. including members of the genera *Pandoravirus*, *Pithovirus*, *Mollivirus*, *Mimivirus* and *Marseillevirus* (Boughalmi et al., 2013; Colson et al., 2013; Legendre et al., 2014; Legendre et al., 2015; Philippe et al., 2013). Although each of these isolates expanded our understanding of the evolutionary history and biological complexity of giant viruses, all are pathogens of *Acanthamoeba* spp., a taxon that is widespread but only representative of a single evolutionary branch of eukaryotes, and which is not a major component in the planktonic communities that dominate the world’s oceans and large lakes.

As knowledge of mimiviruses infecting *Acanthamoeba* spp. has expanded it has become evident based on analysis of metagenomic data that giant viruses and their relatives are widespread and abundant in aquatic systems (Hingamp et al., 2013; Mozar & Claverie, 2014; Schulz et al., 2017). However, except for Cafeteria roenbergensis Virus (CroV) that infects a microzooplankton (Fischer et al., 2010), and the smaller phytoplankton-infecting viruses Phaeocystis globosa virus PgV-16T (Santini et al., 2013) and Aureococcus anophagefferens virus (Moniruzzaman et al., 2014) there are no other members of the *Mimiviridae* that have been isolated and characterized, other than those infecting *Acanthamoeba* spp.

Motivated by the lack of ecologically relevant giant-virus isolates we isolated and screened representative microzooplankton in order to isolate new giant-viruses that can serve as model systems for exploring their biology and function in aquatic ecosystems. Herein, we present Bodo saltans virus (BsV), a giant virus that infects the ecologically important kinetoplastid microzooplankter *Bodo saltans*, a member of the phylum Euglenazoa within the supergroup Excavata. This group of protists is well represented by bodonids in freshwater environments and by diplonemids in the oceans (Flegontova et al., 2016; Simpson, Stevens, & Lukeš, 2006). Kinetoplastids are remarkable for their highly unusual RNA editing and having a single large mitochondrion of circular concatenated DNA (kDNA) that comprises up to 25% of the total genomic content of the cell (Shapiro & Englund, 1995; Simpson et al., 2006), and are well known as causative agents of disease in humans (e.g. Leishmaniasis and sleeping sickness) and livestock (Jackson et al., 2015; Mukherjee, Hodoki, & Nakano, 2015). At 1.39 MB, BsV has the largest described genome within the giant virus family *Mimiviridae*. Based on a recruitment analysis of metagenomic reads, BsV is representative of the most abundant group of giant viruses in the ocean and is the only isolate of the klosneuviruses, a group only known from metagenomic data (Schulz et al., 2017). The BsV genome exhibits evidence of significant genome rearrangements and recent adaptations to its host. Along with *Cafeteria roenbergensis virus* (CroV) and the klosneuviruses, BsV founds a distinct evolutionary group (*Aquavirinae*) that is the dominant clade of *Mimiviridae* within aquatic systems.

## Results

### Isolation and Infection Kinetics

In an effort to isolate giant viruses that infect ecologically relevant organisms, we isolated protistan microzooplankton from a variety of habitats and screened them against their associated virus assemblages. One such screen using water collected from a temperate eutrophic pond in southern British Columbia, Canada, yielded a giant virus that we have classified as Bodo saltans virus, Strain NG1 (BsV-NG1) that infects an isolate of the widely occurring kinetoplastid, *Bodo saltans* (Strain NG, CCCM6296). The addition of BsV to a culture of *Bodo saltans* (~2.5×10^5^ cells ml^−1^) at a virus particle to cell ratio of two, measured by flow cytometry, resulted in free virus particles 18 h later. Viral concentrations peaked at 2.5×10^7^ particles ml^−1^, while host cell density dropped to 25% of uninfected control cultures (Fig. 1A-C). Interestingly, the closely related strain *Bodo saltans* HFCC12 could not be infected by BsV-NG1 suggesting strain specificity.

**Figure 1:**
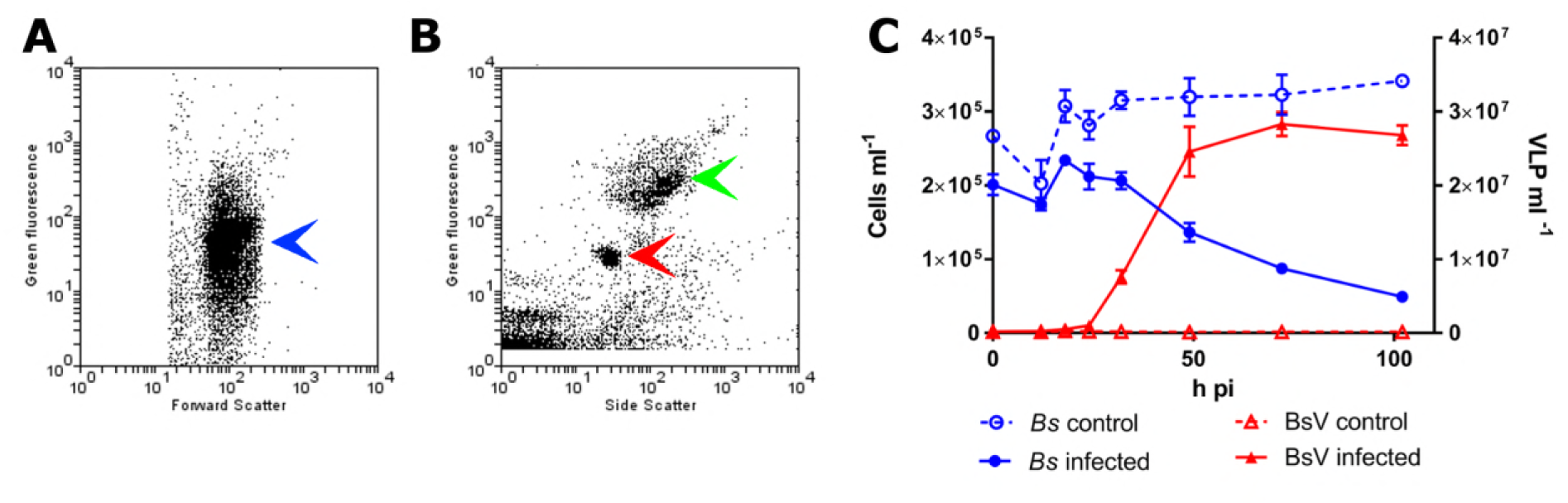
BsV induced lysis observed by flow cytometry. A: Flow cytometry profile of Bodo saltans stained with Lysotracker (blue arrow head) B: Flow cytometry profile of BsV (red arrow head) and bacteria (green arrow head) stained with SYBR Green. C: B. saltans cell and BsV virion (VLP) titer after infection at a PCR of 2

### Virus morphology and Replication Kinetics

Transmission electron microscopy (TEM) revealed that BsV is an icosahedral particle approximately 300 nm in diameter (Fig. 2A). The particle consists of at least six layers akin to observations on Acanthamoeba polyphaga mimivirus (APMV) (Mutsafi, Shimoni, Shimon, & Minsky, 2013). The DNA-containing core of the virion was surrounded by a core wall and an inner membrane, and a putative membrane sitting under a double-capsid layer (Fig. 2A). A halo of approximately 25 nm surrounds the virion. A stargate-like structure that, as observed in APMV, is associated with a depression of the virus core below it and putatively releases the core from the capsid during infection (Fig. 2-Fig.Sup. 1B, D) (Klose et al., 2010; Mutsafi, Fridmann-Sirkis, Milrot, Hevroni, & Minsky, 2014).

**Figure 2:**
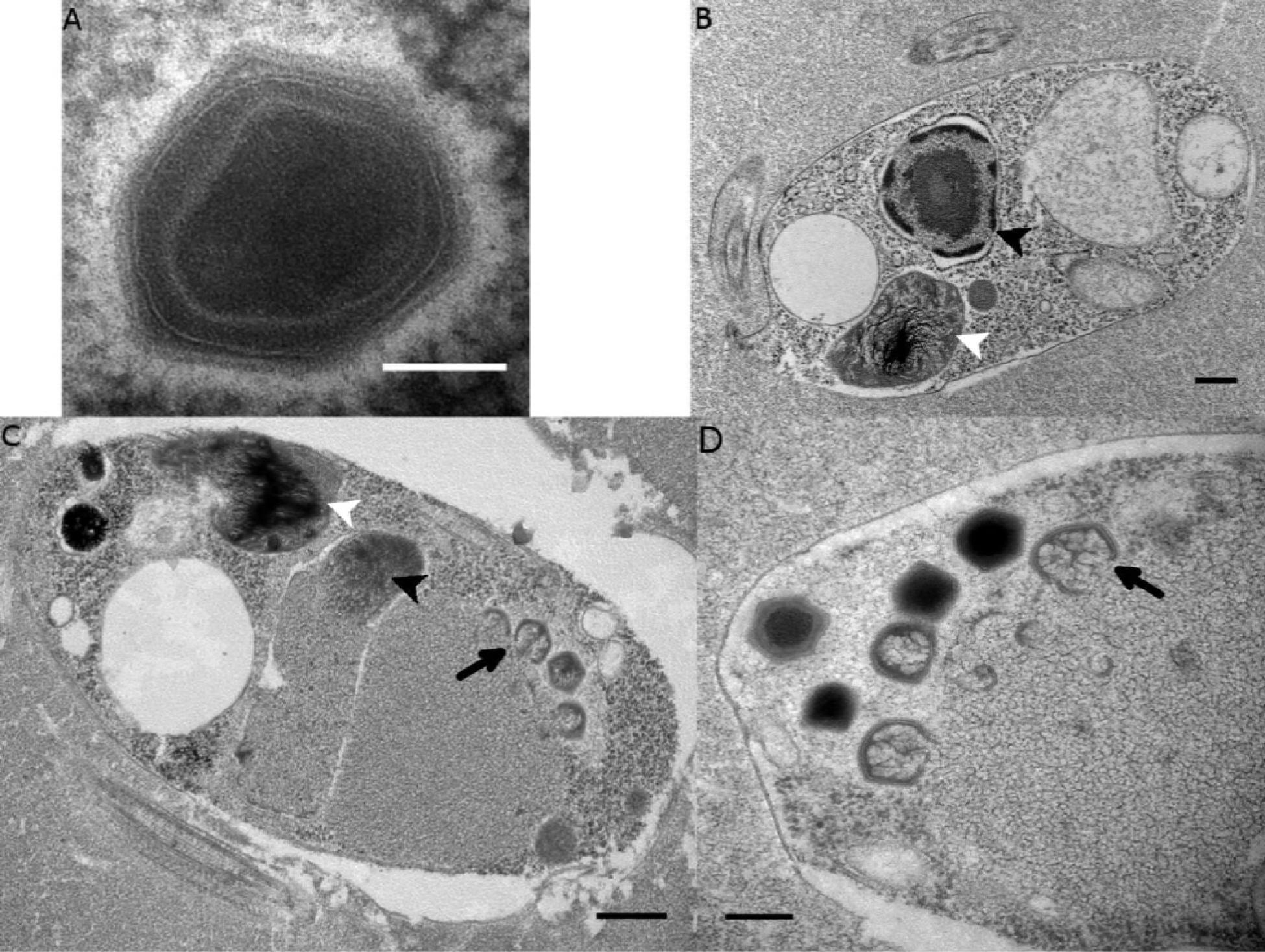
Ultrastructure of BsV particles and replication. A: Mature BsV virion: DNA containing core is surrounded by two putative membranous layers. The capsid consists of at least three additional proteinaceous layers. The bright halo hints to the presence of short (~40nm) fibers as observed in APMV. The top vortex of the virion contains the stargate structure. (Scale bar =100nm) B: Healthy Bodo saltans cell: Nucleus with nucleolus and heterochromatin structures (Back arrow head) and kinetoplast genome (white arrow head) are clearly visible. (Scale bar = 500nm) C: Bodo saltans 24 h post BsV infection: Most subcellular compartments of healthy cells have been displaced by the virus factory now taking up a third of the cell. Virion production is directed towards the periphery of the cell (black arrow). Kinetoplast genome remains intact (white arrow head) while the nuclear genome is degraded (black arrow head; Scale bar = 500nm) D: BsV virion assembly and maturation: Lipid vesicles migrate through the virion factory where capsid proteins attach for the proteinaceous shell. Vesicles burst and accumulate at the virus factory periphery where the capsid assembly completes (black arrow). Once the capsid is assembled, the virion is filled with the genome and detaches from the virus factory. Internal structures develop inside the virion in the cell’s periphery where mature virions accumulate until the host cell bursts. (Scale bar = 500nm)

The healthy *Bodo saltans* cell presents intricate intracellular structures, including the characteristic kinetoplast and a pronounced cytostome and cytopharynx (Fig2B, Fig. 2-Fig.Sup. 1A). In infected cells, virus factories were always observed in the cell’s posterior and particles always matured toward the posterior cell pole in a more spatially organized way compared to other *Mimiviridae* (Fig. 2C, Fig. 2-Fig.Sup. 1C) (Mutsafi, Zauberman, Sabanay, & Minsky, 2010). As infection progressed, the Golgi apparatus disappeared and the nucleus degraded, as evidenced by the loss of the nucleolus and heterochromatin (Fig. 2B, C); yet, the kinetoplast remained intact, as indicated by the persistence of the characteristic kDNA structure (Fig. 1B, C, Fig. 2-Fig.Sup. 1A, C). Virus factories were first observed at 6h post-infection (p.i.) as electron-dense diffuse areas in the cytoplasm. By 12h p.i., the virus factory had expanded significantly and reached a maximum size of about one-third of the host cell, taking up most of the cytoplasm. The first capsid structures appeared at this time. At 18h p.i., the first mature virus particles were observed, coinciding with the first free virus particles observed by flow cytometry (Fig. 1). By 24h p.i., most infected cells were at the late stage of infection with mature virus factories (Fig. 2C, D). During virus replication, membrane vesicles were recruited through the virus factory where capsid proteins accumulated and disrupted the vesicles (Fig. 2D) (Mutsafi et al., 2013). The vesicle/capsid structures accumulated in the periphery of the virus factory where the capsid was formed (Fig. 2C, D). Once the capsid was completed, the viral genome was packaged into the capsid at the vortex opposite to the putative stargate structure (Fig. 2C, D). The internal structures of the virus particle matured in the cell periphery and accumulated below the host cytoplasmic membrane where they often remained for an extended period of time (Fig. 2D, Fig. 2-Fig.Sup. 1D). Besides being released during cell lysis, mature virus particles were observed budding in vesicles from the host membrane, reminiscent of a mechanism described for *Marseillevirus* (Fig. 2-Fig.Sup. 1D) (Arantes et al., 2016).

### Genome organization

Combined PacBio RSII and Illumina MiSeq sequencing resulted in the assembly of a 1,385,869bp linear double-stranded DNA genome (accession number MF782455), making the BsV genome one of the largest complete viral genomes described to date, surpassing those of mimiviruses infecting *Acanthamoeba* spp. The GC content is 25.3% and constant throughout the genome (Fig. 3) and, as reported for other giant viruses, much lower than the ~50% observed for *Bodo spp*. (Jackson et al., 2015; Raoult et al., 2004); this suggests the absence of significant horizontal gene transfer with the host in recent evolutionary history. The genome encodes 1227 predicted open-reading frames (ORFs) with a coding density of 85%, with the ORFs distributed roughly equally between the two strands consistent with the constant GC-skew (Fig. 3). Unlike APMV, BsV does not display a central peak in GC skew and therefore does not have an organized bacterial like origin of replication (Raoult et al., 2004). The genomic periphery has a slightly skewed GC ratio due to the tandem orientation of repeated ORFs. Codon preference is highly biased towards A/T-rich codons and the amino acids Lysine, Asparagine, Isoleucine, and Leucine (10, 9.8, 9.6, 8%), which are preferentially encoded by A/T only triplets. The translation of the predicted ORFs resulted in proteins ranging from 43 to 4840 aa in length with an average length of 320 aa. Promotor analysis revealed a highly-conserved early promotor motif “AAAAATTGA” that is identical to that found in mimiviruses and CroV (Fischer et al., 2010; Priet, Lartigue, Debart, Claverie, & Abergel, 2015). A poorly-conserved late promotor motif “TGCG” surrounded by AT-rich regions was also observed. ORFs are followed by palindromic sequences, suggesting a hairpin-based transcription termination mechanism similar to APMV (Byrne et al., 2009). Several non-coding stretches rich in repetitive sequences were observed, but no function could be attributed to them. Based on a BLASTp analysis, 40% of ORFs had no significant similarity to any other sequences and remained ORFans. Most proteins (27%) matched sequences from eukaryotes; two % of these matched best to sequences from isolates of *B. saltans*. The next largest fraction (22%) were most similar to viruses in the NCLDV group, while the remaining ORFs were most similar to bacterial (9%) or archaeal (1%) sequences. In gene cluster analysis, only 36 % of protein-coding gene clusters are shared with related viruses such as CroV and klosneuviruses, highlighting the low number of conserved core genes amongst these viruses (Fig. 4D). Essential genes for replication, translation, DNA replication and virion structure are located in the central part of the genome, while the periphery is occupied by duplicated genes, including 148 copies of ankyrin-repeat-containing proteins (Fig. 3).

**Figure 3:**
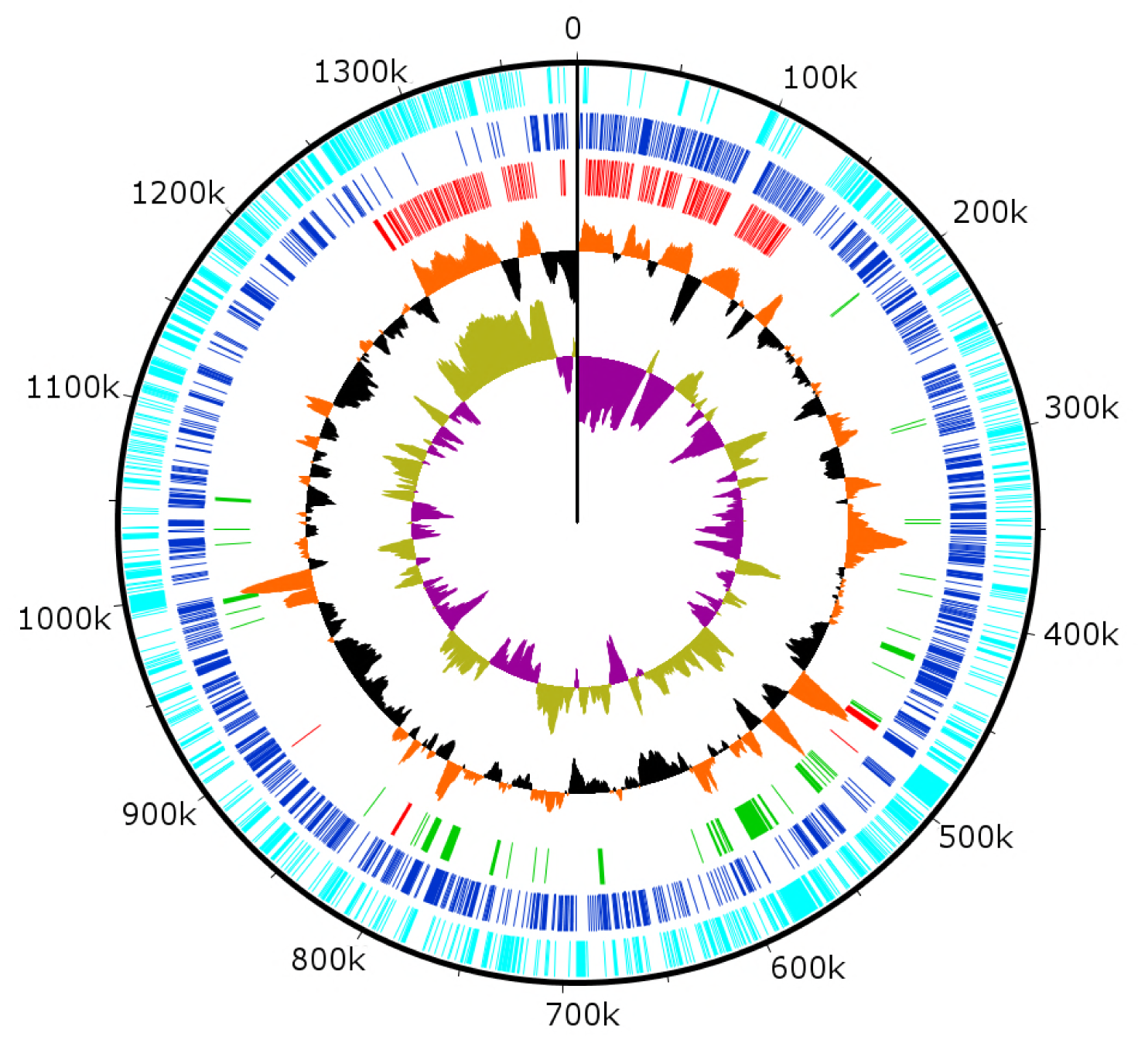
Circularized genome plot of BsV. Circles from inside out: Olive/Purple: GC-skew; Orange/Black: % GC content plotted around average of 25.3%; Red: Ankyrin repeat domain-containing proteins; Green: Essential NCLDV conserved genes; Dark blue: Plus strand encoded ORFs; Light blue: Negative strand encoded ORFs

**Figure 4:**
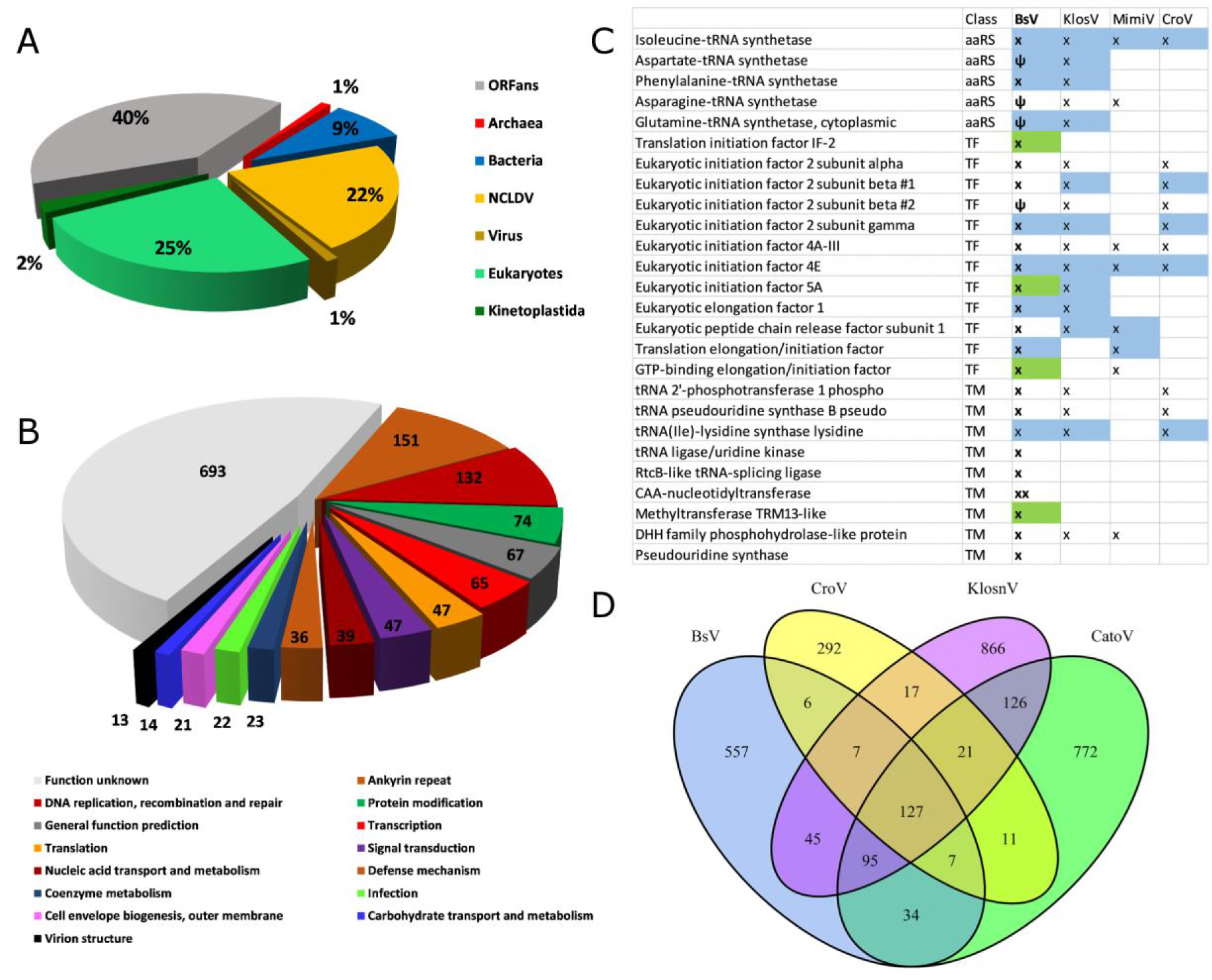
BsV genome content: A: Domain of best BLASTp hits B: Functional assignment of BsV genome content based on BLASTp and CDD rps-BLAST C. BsV Translational machinery: blue-monophyletic group, green: recently host acquired, ψ: pseudogene. A complete list can be found in the Figure Supplement 1 D: Shared gene clusters between BsV, Klosneuvirus (KlosnV), Catovirus (CatoV) and CroV

### Functional genome content

While no function could be attributed to 54% of ORFs, the largest identifiable fraction of annotations are involved in DNA replication and repair (Fig. 4B). Coding sequences for proteins associated with all classes of DNA repair mechanisms were identified including DNA mismatch repair (MutS and Uvr helicase/DDEDDh 3’-5’ exonucleases), nucleotide excision repair (family-2 AP endonucleases), damaged-base excision (uracil-DNA glycosylase and formamidopyrimidine-DNA glycosylase) and photoreactivation (deoxyribodipyrimidine photolyase). The repair pathways are completed by DNA polymerase family X and NAD-dependent DNA ligase. Sequences were also found that putatively code for proteins involved in DNA replication, including several primases, helicases, and an intein-containing family-B DNA polymerase, as well as replication factors A and C, a chromosome segregation ATPase, and topoisomerases 1 (two subunits) and 2. Sequences associated with proteins mediating recombination were also identified including endonucleases and resolvases, as well as the aforementioned DNA repair machinery.

There were 47 sequences identified that matched enzymes involved in protein and signal modification, with the majority being serine/threonine kinases/phosphatases. These are putatively involved in host cell takeover.

The genome of BsV is rich in coding sequences involved in transcription. An early transcription factor putatively recognizing the highly conserved AAAAATTGA motif and a late transcription factor putatively targeting TGCG were identified; whereas, the target sequence of a third transcription factor is unknown. Further, a TATA-binding protein, a transcription initiation factor (TFIIIB) and a transcription elongation factor (TFIIS) were identified that should aid transcription. As well, RNA polymerase subunits a,b,c,e,f,g and I were identified and are assisted by DNA topoisomerases Type 2 and 1B. BsV encodes a putative mRNA specific RNase III, a poly A polymerase, several 5’ capping enzymes and methyl transferases. Transcription is putatively terminated in a manner similar to that described in APMV, as hairpin structures were detected in the 3’ UTR of most putative transcripts (Priet et al., 2015). They are probably recognized and processed by the viral encoded RNase III in a manner similar to APMV (Byrne et al., 2009). After hairpin loop cleavage, the poly-A tail is added by the virally encoded poly-A polymerase. The 5’ capping is accomplished by the virus-encoded mRNA capping enzyme, as well as several capspecific methyltransferases. The extensive cap modification suggests that BsV is independent of the trans-splicing of splice-leader mRNA containing cap structures found in kinetoplastids (Stuart, Allen, Heidmann, & Seiwert, 1997).

BsV also encodes several enzymes associated with nucleic-acid transport and metabolism, including several AT-specific nucleic-acid synthesis pathway components. For instance, adenylosuccinate, thymidylate and pseudouridine synthetases and kinases, as well as ribonucleoside-diphosphate reductase were evident. Other ORFs were associated with nucleotide salvaging pathways, including nucleoside kinases, phosphoribosyl transferases, and cytidine and deoxycytidylate deaminase. A putative mitochondrial carrier protein was identified that, similar to APMV, likely provides dATP and dTTP directly from the kinetoplast to the virus factory, as evident from electron microscopic observations (Monné et al., 2007).

Several genes were identified that are putatively involved in membrane trafficking. a system based on soluble N-ethylmaleimide-sensitive factor (NSF) attachment proteins (SNAPs) and the SNAP receptors (SNAREs) appears to have been acquired from the host by horizontal gene transfer in the recent evolutionary past. In combination with several NSF homologues, including the vesicular-fusion ATPases that also seems to have been acquired from the host. Other proteins putatively involved in membrane trafficking are rab-domain containing proteins, ras-like GTPases, and kinesin motor proteins.

The BsV genome encodes four major capsid proteins that putatively form the outer capsid. One of these proteins contains several large insertions between conserved domains shared among all four capsid proteins, and with 4194 aa boasts a size almost seven times that of its paralogs. This enlarged version of the major capsid protein might be responsible for creating the halo around the virus particles observed by TEM, by producing shortened fibers similar to those observed in APMV (Fig. 2A) (Xiao, 2009). Further, the genome contains two core proteins, several chaperones and glycosylation enzymes suggesting that proteins are highly modified before being incorporated into the virus particle.

There were numerous ORFs that were similar to genes encoding metabolic proteins, like enzymes putatively involved in carbohydrate metabolism. However, no one continuous metabolic pathway could be assembled and therefore these enzymes likely complement host pathways. BsV also encodes coenzyme synthetases such as CoA and NADH and to meet the demand for amino acids that are rare in the host, BsV encodes the key steps in the synthesis pathways of glutamine, histidine, isoleucine, and asparagine.

Another group of genes putatively mediate competitive interactions, either directly with the host, or with other viruses or intracellular pathogens. These include genes involved in the production of several toxins such as a VIP2-like protein as well as putative antitoxins containing BRO domains. Further, a partial bleomycin detox pathway was found, as well as multidrug export pumps and partial restriction modification systems.

While BsV encodes a complex translation machinery, it differs markedly from those described in other NCLDVs. Eukaryotic translation initiation factors include the commonly seen eIF-2a, eIF-2b, eIF-2g, eIF-4A-III and eIF-4E, as well as several pseudogenes related to eIFs. Eukaryotic elongation factor 1 is also present as is eukaryotic peptide chain release factor subunit 1. Notable is the absence of eIF-1; instead, BsV encodes a version of IF-2 that appears to have been acquired from the host and putatively is functionally analogous to eIF-1 in kinetoplastids. The most striking difference to other NCLDVs is the absence of tRNAs. Uniquely among NCLDVs, BsV encodes several tRNA repair genes. These genes include putative RtcB-like RNA-splicing ligase, putative CAA-nucleotidyltransferase, tRNA 2’-phosphotransferase/Ap4A_hydrolase, putative methyltransferase, a TRM13-like protein, pseudouridine synthase and tRNA ligase/uridine kinase. Most of these genes appear to have been recently acquired from the host (Table S1). Other translation modification enzymes found in BsV and other NCLDVs include tRNA(Ile)-lysidine synthase, tRNA pseudouridine synthase B and tRNA 2’-phosphotransferase. Similar to the tRNAs, there are few aminoacyl-tRNA synthetases (aaRS) in BsV. Three of the recognizable aaRS are pseudogenes and show signs of recent nonsense mutations or ORF disruptions by genome rearrangements (aspRS, glnRS, and asnRS). The only complete aaRS proteins are isoleucine-tRNA synthetase, found in all members of the *Mimivridae*, and a phenylalanyl-tRNA synthetase.

### Repeat regions

Genes in the genomic periphery have undergone massive duplication, with 148 copies of ankyrin repeat proteins, mostly present in directional tandem orientation (Fig. 3). These sequences are quite variable and encode between four and 17 ankyrin-repeat domains. There is evidence of very recent sequence duplication resulting in direct or inverted repeat regions that contain complete ankyrin-repeat coding sequences and further expand the repeat clusters (Fig. 3-Fig.Sup. 1A). Interestingly, the 5’ coding region of many ankyrin-repeat containing protein ORFs contain fragments of catalytic domains of essential viral genes such as DNA polymerases or the MutS repair protein (Fig. 3-Fig.Sup. 1C).

### Genomic Mobilome

In contrast to described giant virus genomic mobilomes consisting of virophages and transpovirons, the BsV genome is dominated by inteins, autocatalytic proteinases, and self-splicing group 1 introns (Desnues, 2012; Fischer & Suttle, 2011; Santini et al., 2013; Scola, 2008). These mobile elements target the DNA coding regions of essential genes for virus replication by homing mechanisms deploying unrelated homing endonucleases (Fig. 5). Inteins that are closely related to those in *Mimiviridae* and *Phycodnaviridae* are found in the BsV DNA polymerase family B genes, while unrelated others are found in the DNA dependent RNA polymerase subunits A and B genes (polr2a and polr2b). These RNA polymerase inteins seem to be devoid of an active homing endonuclease and are therefore fixed in these genes, suggesting an evolutionary ancient invasion. The intein in DNA pol B might be an exception, as an HNH endonuclease is located in close genomic proximity and might promote homing in a trans-acting fashion. The group 1 self-splicing introns show signs of at least two independent invasions as the RNA polymerase subunits 1 and 2 genes are occupied by introns with different homing endonucleases (HNH and GIY-YIG type) and different putative ribozymes as evident by their secondary structure (Fig. 5, Fig.Sup. 1B). Subsequently, the homing endonucleases seeded “offspring” introns within the same gene (Fig. 5). These secondary introns show conserved secondary RNA structure, but the lack of the homing endonuclease of their parental intron. Therefore, the secondary introns probably rely on the trans-homing of their parental intron’s endonuclease. The highly conserved sequence for some of the offspring introns (94.4% sequence identity for i1a-i1c, Fig. 5, Fig.Sup. 1A) suggests that these have spread relatively recently, while other introns that only show conservation in their secondary structure probably represent older invasions. Besides proliferating introns, the BsV genome is also home to two distinct actively proliferating transposon classes.

**Figure 5:**
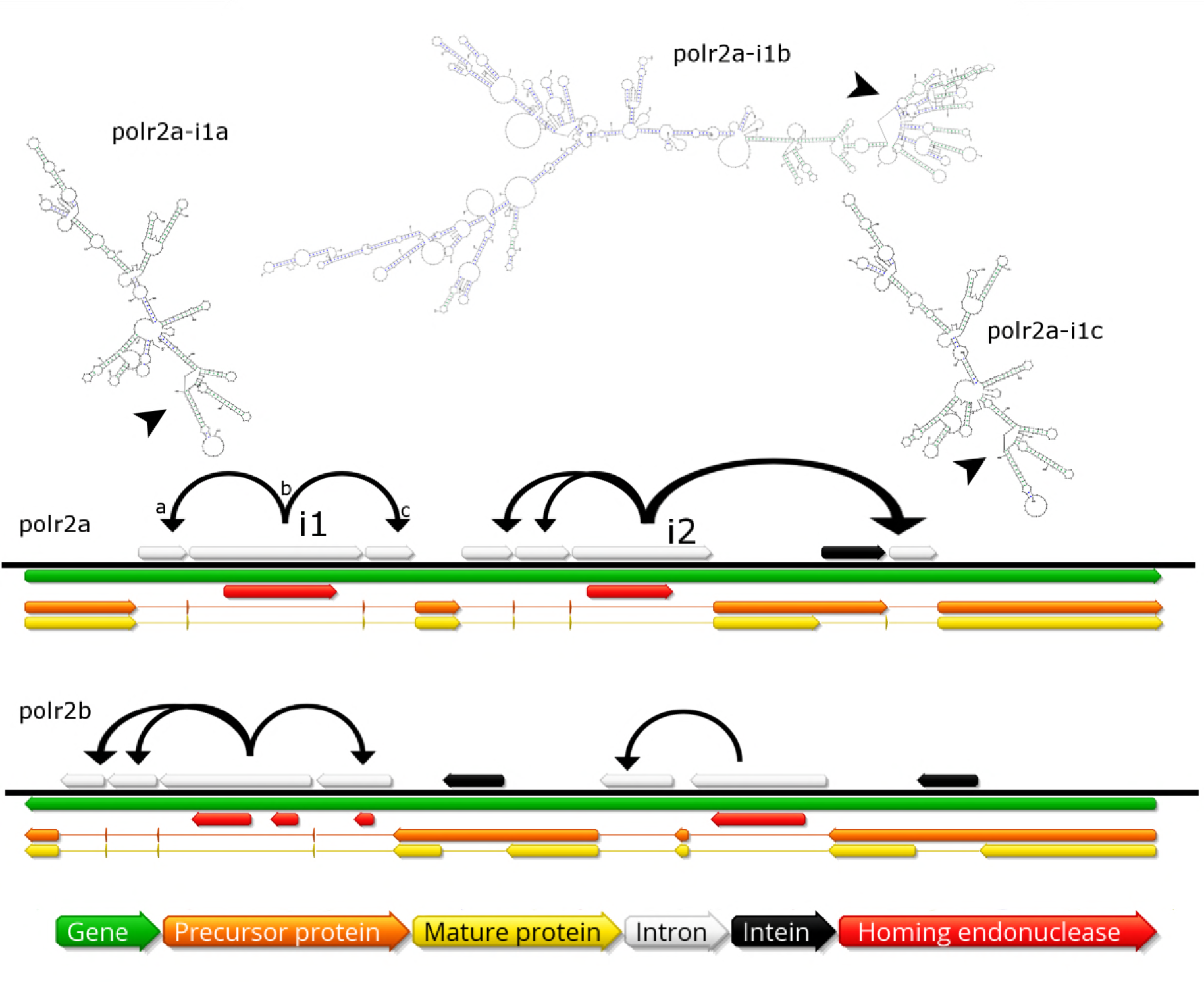
Invasion of RNA polymerase genes by selfish genetic elements. Organization of self-splicing group 1 introns and inteins in genes polr2a and polr2b. Secondary structure of related self-splicing group 1 introns polr2a-i1a through polr2a-i1c shown above the coding sequence with conserved putative self-splicing catalytic site highlighted by arrows. Additional secondary structure predictions and sequence alignment in Supplementary Fig. 3.

### Phylogenetic placement and environmental representation of BsV

Phylogenetic analysis of BsV places it within the *Mimiviridae.* Whole genome phylogenetic analysis based on gene cluster presence/absence of completely sequenced NCLDVs places BsV adjacent to CroV at the intersection of *Acanthamoeba*-infecting large mimiviruses (recently proposed “*Megavirinae”*) and small mimiviruses (“*Mesomimivirinae”*) (Fig. 6 A, B) (Gallot-Lavallee, Blanc, & Claverie, 2017). BsV and CroV are clearly distinct from the *Megavirinae* that show a remarkable amount of specialized genome content not shared with other groups like the *Mesomimivirinae*. Maximum likelihood analysis of five highly conserved NCLDV core genes produced a tree that again resolved three clades within the *Mimiviridae* (Fig. 6C), the *Acanthamoeba*-infecting *Megavirinae*, the *Mesomimivirinae,* and a third, less well-defined clade that includes BsV, CroV, and the klosneuviruses. The longer branch lengths and lower bootstrap support in the third clade, indicate that these viruses are more distantly related to each other than the members of the two established subfamilies; however, they clearly fall outside of the established subfamilies, have overlap in genome content where complete genomes are available (Fig. 4D) and cluster together, suggesting that they are likely a monophyletic group. When metagenomic reads identified to represent the DNA polymerase B of NCLDV from the TARA oceans project were mapped to a DNA polymerase family B tree of the *Mimiviridae*, it was apparent that next to highly represented branches within the *Mesomimivirinae,* the all strains of the new clade represent highly abundant sequences in the environment, far outweighing the *Megavirinae*. This suggests that this clade represents the largest group of identifiable giant viruses in the oceans. Together with the Klosneuviruses and CroV as a distant relative, BsV is critical in defining a new clade of abundant viruses in aquatic environments that are a subfamily within the *Mimiviridae*.

**Figure 6:**
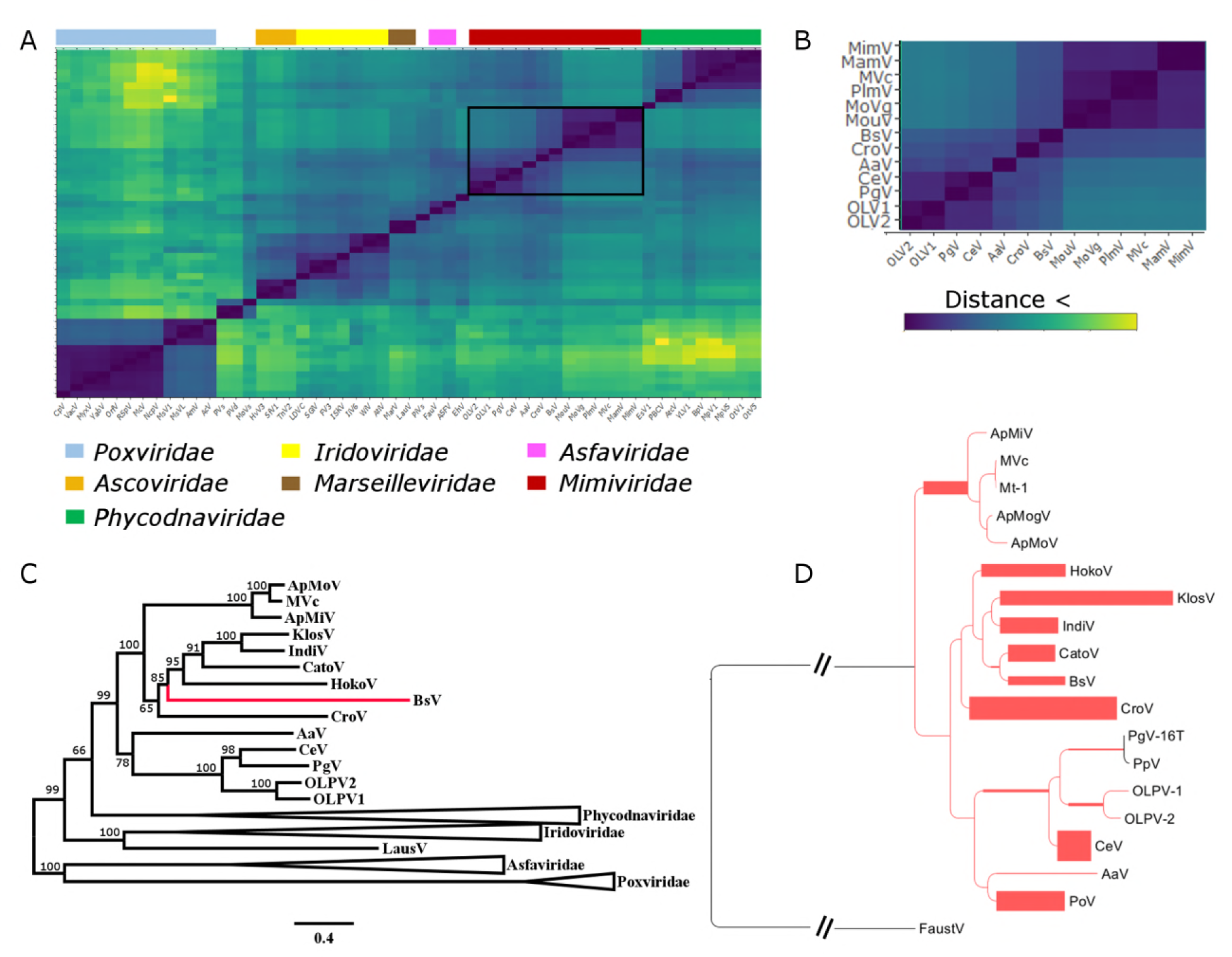
BsV Phylogeny **A:** Phylogenetic distance matrix of complete NCLDV genomes based on whole genome content. Black box highlights the Mimiviridae. **B**: Close-up of the relationships from within the *Mimiviridae* from A. **C**: Maximum likelihood phylogenetic tree of five concatenated NCVOGs from selected NCLDVs. Bootstrap support derived from 1000 bootstrap replicates. Red highlights the position of BsV. D Maximum likelihood phylogenetic tree of DNA Polymerase family B of BsV within the *Mimiviridae*. Branch width correlated to the distribution of 256 TARA ocean NCLDV PolB reads mapped to the tree with pplacer. AaV: *Aureococcus anophagefferens virus*; AcV: *Anomala cumrea entomopoxvirus*; AmV: *Amsacta moorei entomopoxvirus* L; ASFV: *African swine fever virus BA71V*; AtcV: *Acanthocystis turfacea chlorella virus 1*; AtiV: *Aedes taeniorhynchus iridescent virus*; BpV: *Bathycoccus sp. RCC1105 Virus*; BsV: *Bodo saltans virus NG1*; CeV: *Chrysochromulina ericina virus 1B*; CpaV: *Chrysochromulina parva virus BQ1*; CpV: *Canarypox virus*; CroV: *Cafeteria roenbergensis virus BV-PW1*; EhV: *Emiliania huxleyi virus 86*; EsV1: *Ectoparus siliculosus virus 1*; FauV: *Faustovirus E12*; *FsV: Feldmannia species viru*s; FV3: *Frog Virus 3*; HaV: *Heterosigma akashiwo virus 01*; HcV: *Heterocapsa circularisquama DNA virus*; HvV3: *Heliothis virescens ascovirus 3e*; IiV6: *Invertebrate iridescent virus*; InV: *Insectomime virus strain V478*; ISKV: *Infectious spleen and kidney necrosis virus*; LauV: *Lausannevirus*; LDVC: *Lymphocystis disease virus China*; MamV: *Acanthamoeba castellanii mamavirus*; MarV: *Marseillevirus T19*; McV: *Molluscum contagiosum virus Subtype 1*; MimV: *Acanthamoeba polyphaga mimivirus*; MouV: *Acanthamoeba polyphaga moumouvirus*; MoVg: *Moumouvirus goulette*; MoVs: *Mollivirus sibericum*; MpV1: *Micromoas sp. RCC1109 Virus*; MpVS: *Micromoas pusillae Virus SP-1*; MsV1: *Melanoplus sanguinipess entomopoxvirus*; MsVL: *Mythimna separata entomopoxvirus L*; Mt1: *Megavirus terra1*; MVc: *Megavirus chiliensis*; MyxV: *Myxoma virus*; NcpV: *Nile crocodilepox virus*; OLV1: *Organic lake phycodnavirus 1*; OLV2*: Organic lake phycodnavirus 2*; OrfV: *Orf virus*; OtV1: *Ostreococcus tauri virus 1*; OtV5: *Ostreococcus tauri virus 5*; PBCV*: Paramecium bursaria chlorella virus 1*; PgV: *Phaeocystis globosa virus 16T*; PiVs: *Pithovirus sibericum P1084-T*; PlmV: *Powai lake megavirus 1*; PoV: *Pyramimonas orientalis virus*; PpV: *Phaeocystis pouchetii virus*; PVd: *Pandoravirus dulcis*; PVs: *Pandoravirus salinus*; RSpV: *Red squirrel poxvirus UK*; SfV1: *Spodoptera frugiperda ascovirus 1a*; SGiV: *Singapore grouper iridovirus*; TnV2: *Trichoplusia ni ascovirus 2c;* VacV: *Vaccinia virus*; WiV: *Wiseana iridescent virus*; YabV: *Yaba-like disease virus*; YLV1: *Yellowstone lake phycodnavirus 1*

## Discussion

### BsV founds the Aquavririnae, the most abundant subfamily of the Mimiviridae

Particle structure, functional features, like the transcription machinery, and phylogenetic analysis firmly place BsV within the *Mimiviridae*, making it the largest completely sequenced genome of the family. BsV forms a new subfamily with the klosneuviruses and CroV for which we propose the name *Aquavirinae,* clearly distinct from the other subfamilies *Mesomimiviridae* and *Megaviridae* (Fig. 6). The high representation of the *Aquavirinae* in metagenomic reads suggests that they represent the largest group of giant viruses in the oceans (Hingamp et al., 2013). The detection of Klosneuviruses in low complexity fresh water metagenomes further supports the global prevalence of the *Aquavirinae*.

### BsV has acquired a host mechanism to facilitate membrane fusion, employed during infection and virion morphogenesis

The SNAP/SNARE membrane fusion system found in BsV appears to have been recently acquired from the bodonid host via horizontal gene transfer. This system could mediate membrane fusion in a pH-dependent manner (Itakura, Kishi-Itakura, & Mizushima, 2012). Accordingly, we propose a phagocytosis based infection strategy for BsV: As described for APMV, BsV is ingested through the cytostome and is phagocytosed in the cytopharynx before being transported in a phagosome towards the posterior of the cell (Mutsafi et al., 2010); here the viral SNAP/SNARE interacts with the host counterparts to initiate the fusion of the inner virus membrane with the phagolysosomal membrane upon phagolysosome acidification, releasing the viral genome into the cytoplasm. This scenario is supported by the localization of the virus factory at the posterior of the cell and virus particle structure (Fig. 2A,C and Fig. 2 Fig.Sup. 1B-C). According to this hypothesis, SNAP/SNARE proteins must be present membranes of the mature virus particles and only get exposed after the stargate opens. The SNAP/SNARE system might also be involved in recruiting membrane vesicles from host organelles to the virus factory during maturation of the virus particle as has been described for pox viruses (Fig. 2D) (Laidlaw et al., 1998).

### The BsV possesses an arsenal of mechanisms to fend of competitors

As a representative of environmentally highly abundant viruses, BsV might regularly experience competition for host resources. The putative toxin-antitoxin systems observed in BsV might be involved in competing with other parasites of viral or prokaryotic nature for these resources, by inhibiting their metabolism or damaging their genome as proposed for APMV (Boyer et al., 2011). Most remarkable, however, are the site-specific homing endonucleases encoded by the self-splicing group 1 introns and inteins that have invaded several genes essential for BsV replication. These invasions also seem to be part of the competitive arsenal of BsV, vending off related virus strains competing for abundant and common hosts such as bodonids. During superinfection of two related viruses, having selfish elements encoding homing endonucleases targeting essential genes, the home of the selfish element in the original virus, might be a competitive advantage. As the two competing virus factories are established in the cytoplasm, the endonucleases encoded by the intron or intein cleave the unoccupied locus in the genome of the intron/intein free virus. Thus, the intron-containing virus is reducing the ability of the competing virus to replicate (Fig. 7). A similar mechanism has been described for competing phages in which an intron-encoded or derived homing endonuclease mediates marker exclusion during superinfection, causing selective sweeps of genes in the vicinity of the endonuclease through the phage population (Belle, Landthaler, & Shub, 2002; Goodrich-Blair & Shub, 1996; Kutter et al., 1995). More credence to this hypothesis is given by the RNA polymerase sequences encoded by the proposed catovirus and klosneuvirus (Schulz et al., 2017). These genes (polr2a and polr2b) are fragmented in a manner similar to that observed in BsV, and also appear to encode homing endonucleases between gene fragments, suggesting the presence of self-splicing introns is common in the *Aquavirinae*. Since the hosts for klosneuviruses are unknown, it is possible that they compete with BsV for the same hosts, or at least experience similar competition. The presence of non-fixed inteins in other giant viruses hints to past invasions and selection for inteins in a manner resembling the hypothesis proposed here for introns (Culley, Asuncion, & Steward, 2009; Gallot-Lavallee et al., 2017). The retention of fixed inteins in BsV and other giant viruses suggests that there are still viruses in the environment that encode the relevant endonucleases that apply selective pressure to retain the inteins. A similar situation might explain the presence of an intein in the DNA polymerase of *Pandoravirus salinus*, but not in *P. dulcis*. Thus, pandoraviruses may be an excellent model to experimentally explore the proposed mechanism of intron homing endonuclease mediated competition.

**Figure 7:**
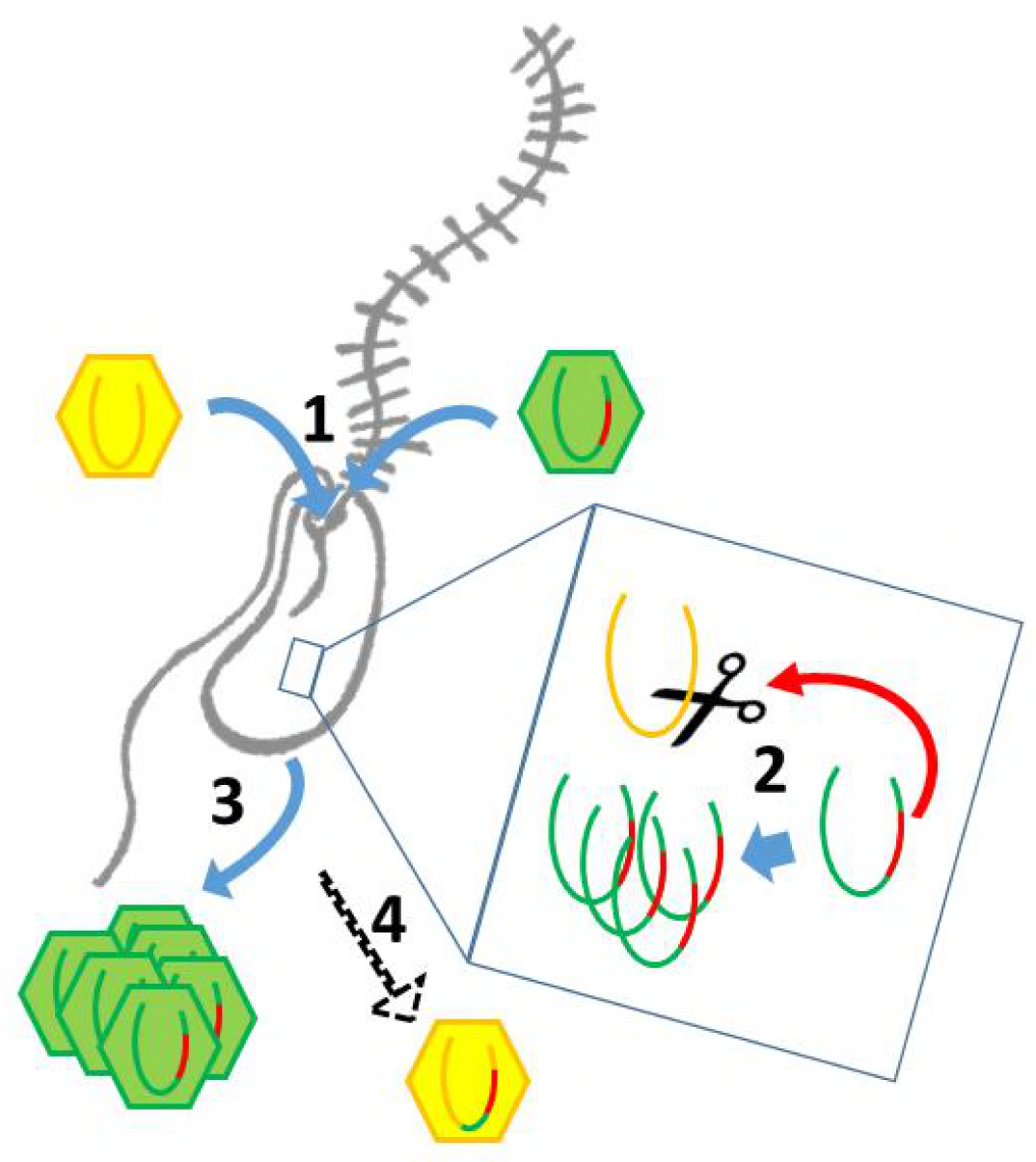
Intron/intein encoded endonuclease mediated interference completion between related viruses: 1) two related viruses infect the same host cell. The green virus genome contains a selfish element encoding a homing endonuclease. 2) During initial replication, the endonuclease is expressed and cleaves the unoccupied locus on the yellow virus’ genome impairing its replication. 3) Due to suppressing its competitor’s replication, the majority of the viral progeny is of the green virus’ type. 4) The yellow virus can rescue its genome by using the green virus’ genome as a template. This creates a chimeric genome containing the selfish element and the endonuclease as well as adjacent sequences originating from the green virus’ genome.

### The translation machinery of giant viruses is a homoplasic trait

The absence of tRNAs in the BsV genome is remarkable since tRNAs are found in all genomes of giant viruses and even in many moderately sized NCLDV genomes. This might be an adaptation to the unusual RNA modification found in kinetoplastids that also encompasses tRNA editing (Alfonzo & Lukeš, 2011; Stuart et al., 1997). BsV likely cannot replicate this unusual editing, and thus relies on using host tRNAs. Hence, BsV encodes tRNA repair genes to compensate for the lack of tRNA synthesis and to maintain the available tRNA pool in the host cell. Most of these genes appear to have been recently acquired from the host (Table S1). Like the tRNAs, most virus-encoded aminoacyl-tRNA synthetases might not be able to recognize the highly modified tRNAs present in the host and are therefore degrading in the absence of positive selective pressure (Fig. 4, Fig.Sup. 1). The loss and resulting heterogeneity in translational machinery compared to klosneuviruses, combined with the apparent diverse origin of these genes, suggests that the translation machinery found in giant viruses is the result of rapid adaptation by gene acquisition via horizontal gene transfer, as has been recently proposed by Shultz et al. (Schulz et al., 2017). Furthermore, BsV demonstrates that such genes can be readily purged from the virus genome if they not required in a new host. Thus, BsV provides further evidence that the translation machinery encoded by NCLDVs is a homoplasic trait, need not be ancient in origin, and is not evidence of a fourth domain of life.

### An inflated genomic accordion due to evolutionary arms races is responsible for genome gigantism in BsV and sheds light on giant virus evolution

The 148 copies of ankyrin-repeat domain proteins in the genomic periphery of BsV are telltale signs of an expanded genomic accordion (Fig. 3) (Elde et al., 2012). The observation of almost identical sequences in the very periphery of the genome is consistent with the genomic accordion hypothesis, in which the most recent duplications are closest to the genome ends (Fig. 3, Fig.Sup. 1A,B). The genomic recombinations causing the gene duplications can also lead to the disruption of coding sequences that might explain the comparatively low coding density of BsV. Proteins with ankyrin-repeat domains are multifunctional attachment proteins that in pox viruses determine host range by inhibiting host innate immune system functions (Camus-Bouclainville et al., 2004). The presence of fragments of the catalytic domains of essential viral genes in many ankyrin-repeat containing genes is of further importance (Fig. 3, Fig.Sup.1 C). This suggests a decoy defense mechanism, where these fusion proteins mimic the targets of host anti-viral defense systems disrupting essential viral functions. By acting as decoy targets, they immobilize the proposed host factors upon binding via their ankyrin-repeat domains similar to what occurs in vaccinia virus (Elde, Child, Geballe, & Malik, 2009). The immobilized host factors might even be degraded in a ubiquitin dependent manner reminiscent of the situation in pox viruses as suggested by the presence of several ubiquitin conjugating enzymes encoded in the BsV genome (Sonnberg, Seet, Pawson, Fleming, & Mercer, 2008). An ankyrin-repeat based defense system might explain the observation of cells surviving or avoiding infection that can persist in the presence of the virus (Fig. 1C). Alternatively, the protein-protein interaction mode of ankyrin repeat proteins might aid attachment and induction of phagocytosis as the bodonid host cells have changing surface antigens (Jackson et al., 2015). Whatever the true function of the ankyrin-repeat proteins might be, they clearly highlight the importance of the genomic accordion in giant virus genome evolution driven by evolutionary arms races and complement previous observations of a contracting genomic accordion in APMV (Boyer et al., 2011).

### Summary

Bodo saltans virus has the largest genome size of any member of the *Mimiviridae* isolated to date, despite not infecting the usual amoeboid host of previously known truly giant viruses. BsV highlights the genomic plasticity of giant viruses via the genomic accordion that employs large-scale genome expansions and contractions via non-homologous recombination. The recent duplications in BsV demonstrate genome expansion in action and exemplify the mechanisms leading to genome gigantism in the *Mimiviridae*. Further, the putative function of the expanding genes suggests strong evolutionary pressures placed on these viruses by virus-host arms races as the driver of genomic expansions. The genomic plasticity is further apparent in the translational machinery that shows signs of recent gene loss and rapid adaptation to its bodonid host. This emphasizes that the translational machinery of giant viruses is indeed an acquired homoplasic trait not derived from a common ancestor. An invasion of selfish elements in essential genes suggests interference competition among related viruses for shared hosts. BsV defines a new subfamily of the *Mimiviridae* that based on metagenomic data represents the most abundant giant viruses in aquatic ecosystems. Further, it is the first DNA virus isolate described that infects the eukaryotic supergroup Excavata, a major evolutionary lineage. Bodo saltans virus provides significant new insights into giant viruses and their biology.

## Materials and methods

### Sampling and Isolation

Virus concentrates were collected from 11 fresh water locations in southern British Columbia, Canada. To concentrate giant viruses, 20 liter water samples were prefiltered with a GF-A filter (Millipore, Bedford, MA, USA; nominal pore size 1.1um) over a 0.8um PES membrane (Sterlitech, Kent, WA, USA). Filtrates from all locations were pooled and were concentrated using a 30kDa MW cut-off tangential flow filtration cartridge (Millipore, Bedford, MA, USA) (Curtis A Suttle, Chan, & Cottrell, 1991).

The BsV host organism*, Bodo saltans* NG was isolated from a near sediment water sample of a pond in Nitobe Gardens at the University of British Columbia Canada (49°15’58"N, 123°15’34"W) and clonal cultures after end point dilutions were maintained in modified DY-V artificial fresh water media with yeast extract and wheat grain. The identity of the host organism was established by 18S sequencing and the strain was deposited at the Canadian Center for the Culture of Microorganisms (http://www3.botany.ubc.ca/cccm/) reference number CCCM 6296 (von der Heyden & Cavalier-Smith, 2005). *Bodo saltans* cultures were inoculated at approximately 2×10^5^ cells/ml with the giant virus concentrate. Cell numbers were screened by flow cytometry compared to a mock-infected control culture (LysoTracker Green (Molecular Probes) vs. FSC on FACScalibur (Becton-Dickinson, Franklin Lakes, New Jersey, USA))(Rose, Caron, Sieracki, & Poulton, 2004). After a lytic event was observed, the lysate was filtered through a 0.8um PES membrane (Sterlitech) to remove host cells. The lytic agent was propagated and a monoclonal stock was created by three consecutive end point dilutions. The concentrations of the lytic agent were screened by flow cytometry using SYBR Green (Invitrogen Carlsbad, California, USA) nucleic acid stain after 2% glutaraldehyde fixation (vs SSC) (Brussaard, 2004). The flow cytometry profile was presented as a tight population of homogeneous particles clearly distinct from bacteria, phage, and eukaryotes. The similarity to the flow cytometry profile of Cafeteria roenbergensis virus suggested that the lytic agent was in deed a giant virus.

### Transmission electron microscopy

#### Negative staining

Bodo saltans lysates after BsV infection were applied to the carbon side to a formvar-carbon coated 400 mesh copper grid (TedPella, CA, USA) and incubated at 4°C in the dark overnight in the presence of high humidity. Next, the lysate was removed and the grids were stained with 1% Uranyl acetate for 30 seconds before observation on a Hitachi H7600 transmission electron microscope at 80 kV.

#### High-pressure freezing and ultra-thin sectioning

Exponentially growing *B. saltans* cultures were infected at a concentration of 5×10^5^ cells ml^−1^ with BsV at a relative particle to cell ratio of ~5 to ensure synchronous infection. Cells were harvested from infected cultures at different time points (6, 12, 18, 24 h post infection) as well as from uninfected control cultures. Cells from 50 ml were pelleted in two consecutive 10 min at 5000 xg centrifugation runs in a Beckmann tabletop centrifuge. Pellets were resuspended in 10-15 µl DY-V culture medium with 20% (w/v) BSA and immediately place on ice. Cell suspensions were cryo-preserved using a Leica EM HPM100 high-pressure freezer. Vitrified samples were freeze-substituted in a Leica AFS system for 2 days at −85°C in a 0.5% glutaraldehyde 0.1% tannic acid solution in acetone, then rinsed ten times in 100% acetone at −85°C, and transferred to 1% osmium tetroxide, 0.1% uranyl acetate in acetone and stored for an additional 2 days at −85°C. The samples were then warmed to −20°C over 10 hours, held at −20°C for 6 hours to facilitate osmication, and then warmed to 4°C over 12 hours. The samples were then rinsed in 100% acetone 3X at room temperature and gradually infiltrated with an equal part mixture of Spurr’s and Gembed embedding media. Samples were polymerized in a 60°C oven overnight. 50 nm thin sections were prepared using a Diatome ultra 45° knife (Diatome, Switzerland) on an ultra-microtome. The sections were collected on a 40x copper grid and stained for 10 min in 2% aqueous uranyl acetate and 5 min in Reynold’s lead citrate. Image data were recorded on a Hitachi H7600 transmission electron microscope at 80 kV. Image J was used to compile all TEM images. Adjustments to contrast and brightness levels were applied equally to all parts of the image.

### Virus concentration and sequencing

For Illumina sequencing, exponentially growing *B. saltans* cultures were infected at a concentration of approximately 5×10^5^ cells ml^−−1^ with BsV lysate (10^7^ VLP ml^−1^) at a multiplicity of infection (MOI) of ~0.5. After four days, when host cell densities had dropped below 30%, cultures were centrifuged in a Sorvall SLC-6000 for 20 min, 5000 rpm, 4°C to remove remaining host cells and the supernatant was consecutively subjected to tangential flow filtration with at 30kDa cut-off (Vivaflow PES) and concentrated approximately 100x. Viral concentrates were subjected to ultracentrifugation at 28,000 rpm, 15°C for 8h in a Ti90 fixed angle rotor (Beckman-Coulter, Brea, California, USA). Pellets were resuspended and virions lysed using laurosyl acid and proteinase K subjected to pulsed-field gel electrophoresis on a CHEF II pulse field gel electrophoresis aperture (BioRad) for 25h at 14°C in a 0.8% LMP agarose gel with 60-180S switchtimes and 16.170 ramping factor in 0.5 TBE under 5.5V/cm and 120°. Genomic DNA was visualized under UV light after 30min SYBR gold (Invitrogen Carlsbad, California, USA) staining. The dominant PFGE band belonging to genomic BsV DNA (1.35Mb) was cut and DNA was extracted using a GELase kit (Illumina, San Diego) and ethanol purified according to manufacturer’s protocol. Libraries were prepared using the Illumina Nextera XT kit (Illumina, San Diego, CA, USA) as per manufacturer’s recommendation and library quality and quantity were checked by Bioanalyzer 2100 with the HS DNA kit (Agilent Technology). 300bp paired-end sequencing was performed on an Illumina MiSeq platform by UCLA’s Genoseq center (Los Angeles, CA, USA) to a nominal sequencing depth of 800x. Sequence quality was examined using FastQC (http://www.bioinformatics.bbsrc.ac.uk/projects/fastqc/) and sequence reads were quality trimmed (parameters: minlen=50 qtrim=rl trimq=15 ktrim=r k=21 mink=8 ref=$adapters hdist=2 hdist2=1 tbo=t tpe=t) and cleared of human (parameters: minid=0.95 maxindel=3 bwr=0.16 bw=12 quickmatch fast minhits=2 qtrim=lr trimq=10) and PhiX (parameters: k=31 hdist=1) sequences against the whole respective genomes using BBMap v35 (http://sourceforge.net/projects/bbmap/).

For PacBio sequencing, BsV was concentrated using precentrifugation and TFF analogously to the Illumina sequencing step. Next, the concentrate was further concentrated by sedimenting it onto a 40% Optiprep 50 mM Tris-Cl, pH 8.0, 2mM MgCl_2_ cushion for 30 min at 28,000 rpm, 15°C in a SW40Ti rotor in an ultracentrifuge (Beckman-Coulter, Brea, California, USA). An Optiprep (Sigma) gradient was created by underlaying a 10% Optiprep solution in 50 mM Tris-Cl, pH 8.0, 2mM MgCl_2_ with a 30% solution followed by a 50% solution and was equilibration over night at 4°C. One ml of viral concentrate from the 40% cushion was added atop the gradient and the concentrate was fractionated by ultracentrifugation in an SW40 rotor for 4h at 25000rpm and 18°C. The viral fraction was extracted from the gradient with a syringe and washed twice with 50 mM Tris-Cl, pH 8.0, 2mM MgCl_2_ followed by centrifugation in an SW40 rotor for 20 min at 7200rpm and 18°C and were finally collected by centrifugation in an SW40 rotor for 30 min at 7800rpm and 18°C. Purity of the concentrate was verified by flow cytometry (SYBR Green (Invitrogen Carlsbad, California, USA) vs SSC on a FACScalibur (Becton-Dickinson, Franklin Lakes, New Jersey, USA). High molecular weight genomic DNA was extracted using phenol-chloroform-chloroform extraction. Length and purity were confirmed by gel electrophoresis and Bioanalyzer 2100 wit the HS DNA kit (Agilent Technology). PacBio RSII 20kb sequencing was performed by the sequencing center of the University of Delaware. Reads were assembled using PacBio HGAP3 software with 20 kb seed reads and resulted in a single viral contig of 1,384,624bp, 286.1x coverage, 99.99% called bases and a consensus concordance of 99.9551% (Chin et al., 2013).

Cleaned up Illumina reads were mapped to the PacBio contig to confirm the PacBio assembly as well as extending the contig’s 5’ end by 1245bp to a total viral genome length of 1,385,869bp.

### Annotation

Open reading frames were predicted using GLIMMER with a custom start codon frequency of ATG, GTG, TTG, ATA, ATT at 0.8,0.05,0.05,0.05,0.05 as well as stop codons TAG, TGA, TAA, minimum length 100bp, max overlap 25, max threshold 30.

Promoter motives were analyzed by screening the 100bp upstream region of CDS using MEME. tRNAs were predicted with tRNAscanSE (Lowe & Eddy, 1997). Group 1 introns were predicted by disruptions in coding sequences and secondary RNA structure was predicted using S-fold(Ding, Chan, & Lawrence, 2004). Intron splicing was confirmed using RT-PCR and Sanger sequencing with gene-specific primers designed to span the predicted splice sited predicted by S-fold. Functional analysis of CDS was performed after translation with BLASTp against the nr database with an e-value threshold of 10^−5^ as well as rps conserved domain search against CDD v3.15. Coding sequences were manually assigned to functional classes based on predicted gene function using Geneious R9 (Kearse et al., 2012). The annotated genome of BsV-NG1 was deposited in GenBank under the accession number MF782455. The *Bodo saltans* NG 18S sequence was deposited in GenBank under the accession number MF962814.

### Phylogenetics

Whole genome content phylogeny was performed by OrthoMCL (Li, Stoeckert, & Roos, 2003). Available whole genome sequences of NCLDV from NCBI were downloaded. We first performed gene clustering using OrthoMCL (42) with standard parameters (Blast E-value cutoff = 10^−5^ and mcl inflation factor = 1.5) on all protein-coding genes of length ≥ 100 aa. This resulted in the definition of 3,001 distinct clusters. We computed a presence/absence matrix based on the genes clusters and calculated a distance matrix using the according to Yutin et al. 2009 (Yutin, Wolf, Raoult, & Koonin, 2009).

Alignments of aa sequences were performed in Geneious R9 using MUSCLE with default parameters (Edgar, 2004). Proteins used for the concatenated NCVOG tree were DNA polymerase elongation subunit family B (NCVOG0038), D5-like helicase-primase (NCVOG0023), packaging ATPase (NCVOG0249), Poxvirus Late Transcription Factor VLTF3-like (NCVOG0262), and DNA or RNA helicases of superfamily II (NCVOG0076) (Yutin, Wolf, & Koonin, 2014). Residues not present in at least 2/3 of the sequences were trimmed and ProtTest 3.2 was used for amino-acid substitution model selection (Darriba, Taboada, Doallo, & Posada, 2011). Maximum likelihood trees were constructed with RAxML rapid bootstrapping and ML search with 1000 Bootstraps utilizing the best fitting substitution matrixes determined by protest (Stamatakis, 2014). Maximum likelihood trees of translational genes, based on alignments by Schulz et al. 2017 where available, were constructed using PhyML (Guindon et al., 2010; Schulz et al., 2017). Trees were visualized in Figtree (A. Rambaut - http://tree.bio.ed.ac.uk/software/figtree/).

Translated environmental assemblies identified by Hingamp et al. as representing NCLDV DNA polymerase B family genes were mapped to a *Mimiviridae* DNA polymerase B family reference tree created as described above with pplacer (Hingamp et al., 2013; Matsen, Kodner, & Armbrust, 2010). Environmental reads were aligned to the reference alignment using clustalw and were mapped under Bayesian setting. The fat tree was visualized with Archaeopteryx (https://sites.google.com/site/cmzmasek/home/software/archaeopteryx). Of the 401 input sequences, 256 mapped within the *Mimiviridae* and are displayed in the figure.

## Acknowledgments

We thank members of the Suttle Lab, past and present, and especially Andrew Lang and Matthias Fischer for helpful comments. As well, the assistance of Jan Finke, Marli Vlok, Marie-Claire Veilleux-Foppiano and Amy M. Chan is greatly appreciated, who directly assisted in sample collection, laboratory work, and ensuring the equipment and supplies were available to do the work. Further, we would like to thank Grieg Steward for providing insights into sample preparation. Denis Tikhonenkov generously provided a culture of *Bodo saltans* HFCC12. John Archibald and Julius Lukeš provided valuable feedback on bodonid biology. Thor Veen assisted with R script writing. The work was supported by grants to Curtis A. Suttle from the Natural Sciences and Engineering Research Council of Canada (NSERC; 05896), Canadian Foundation for Innovation (25412), British Columbia Knowledge Development Fund, and the Canadian Institute for Advanced Research (IMB). Christoph Deeg was supported by a fellowship from the German Academic Exchange Service (DAAD) and Cheryl E.T. Chow by a Centre for Microbial Diversity and Evolution postdoctoral scholarship funded through the Tula Foundation.

## Supplemental Figures

**Figure 2 –Figure Supplement 1:**
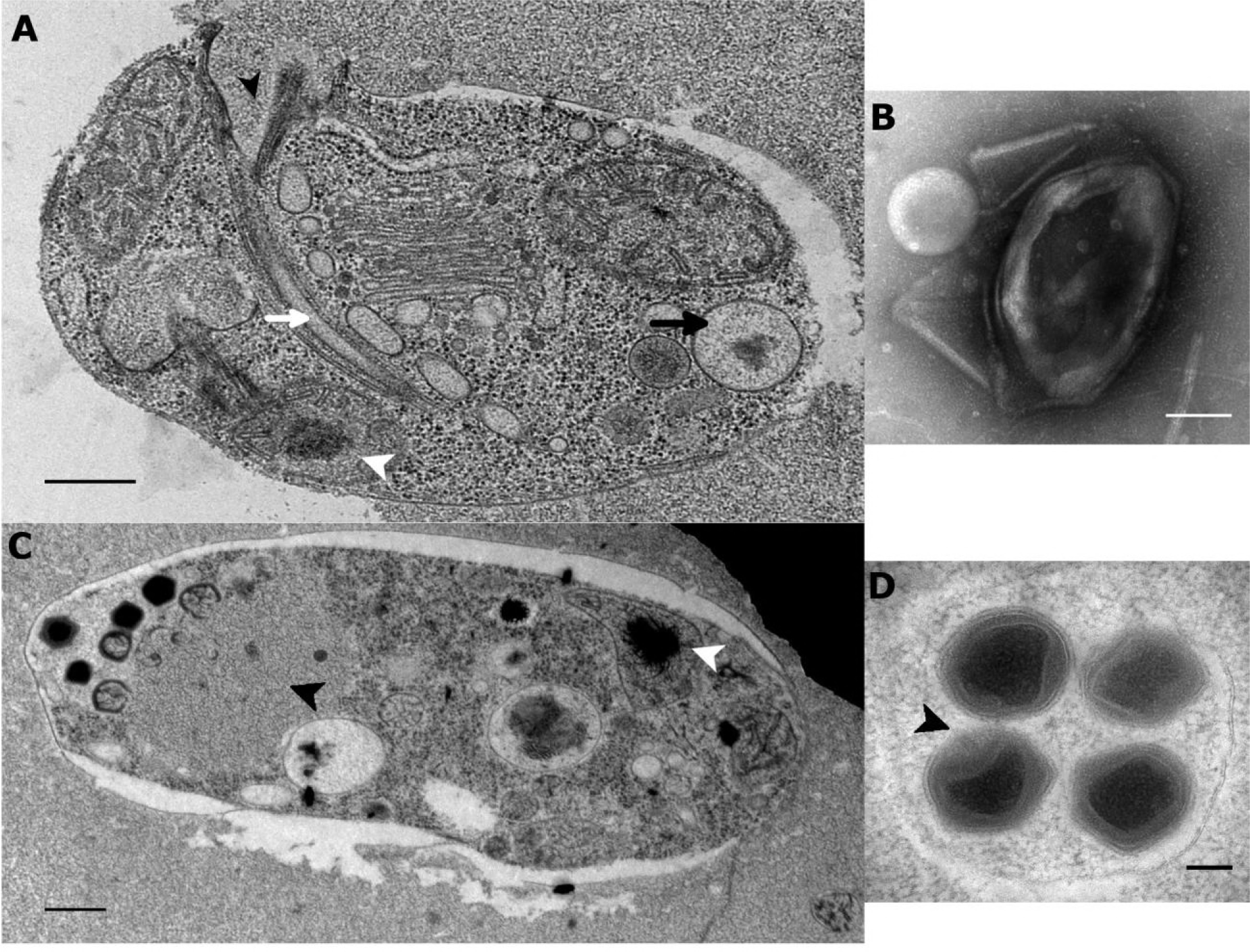
Ultrastructure of BsV particles and replication. A: Healthy Bodo saltans cell: Visible structures include the cytostome (black arrow head), the Golgi, mitochondrial arms protruding from the kinetoplast center with the kinetoplast genome (white arrow head), the flagellar root of both flagella as well as several vacuoles (back arrow: food vacuole containing partially digested bacterial prey) are visible traveling from the cytopharynx (white arrow) to the posterior cell pole (Scale bar = 500nm) B: Negative staining of a BsV particle with a blossom like opened stargate C: Bodo saltans cell 24h post BsV infection showing degraded intracellular structures and extensive BsV virion factory (black arrow head). D: BsV virions inside a vesicle. A closed stargate is visible at the apex of the bottom left virion (black arrow head).

**Figure 3 –Figure Supplement 1:**
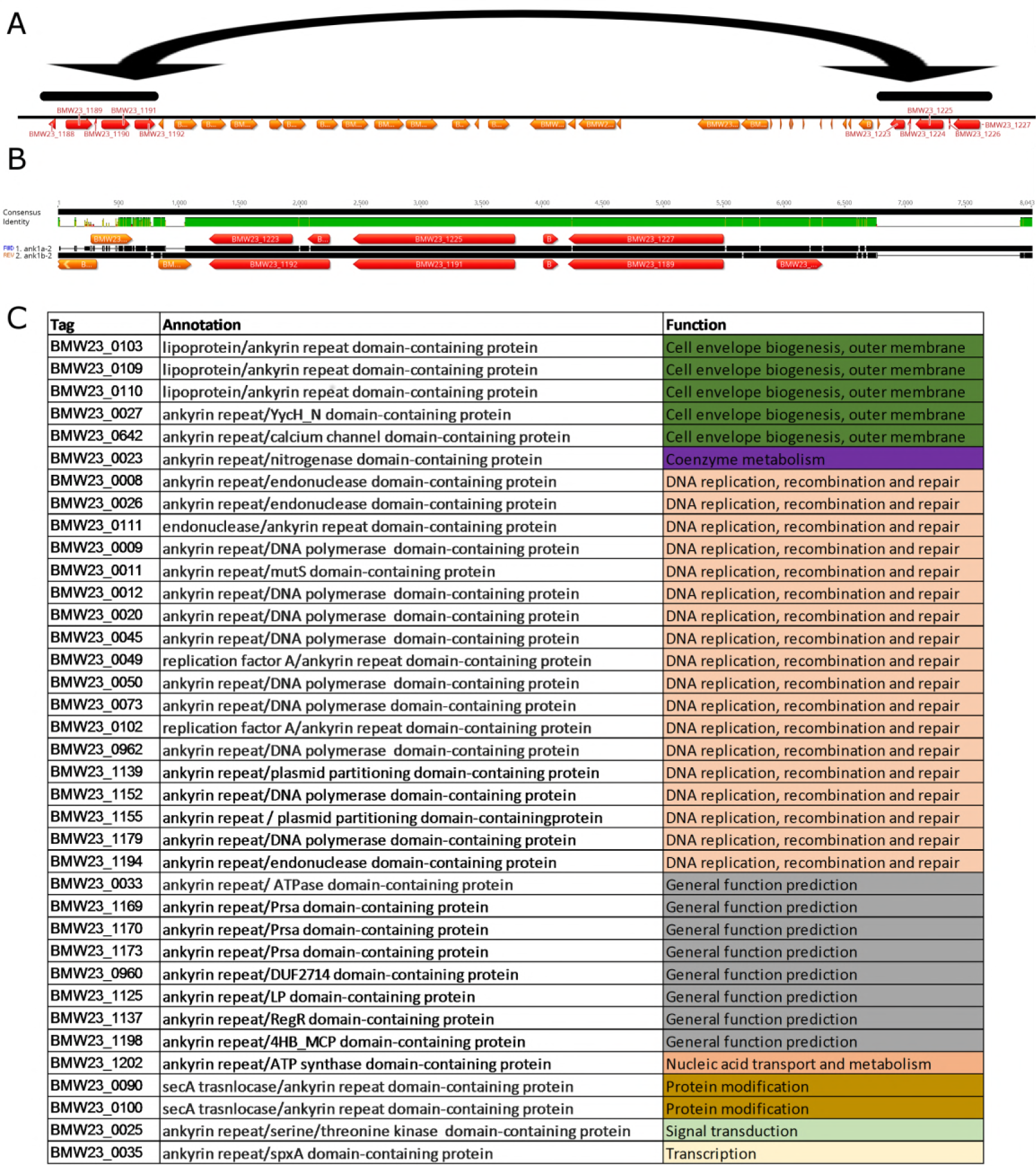
Ankyrin repeat domain-containing proteins: A: Location of recently duplicated region in the 3’ end of the genome (highlighted in black bars, duplicated CDS in red) B: Alignment of recently duplicated sequence from A, encoding several Ankyrin repeat domain-containing proteins. A region containing four CDS was duplicated in a reverse complement orientation maintaining the integrity and sequence identity of the original sequence (% DNA ID in the central region is 99.7%) C: ID, annotation and functional class of ankyrin repeat domain-containing proteins with a recognizable domain fragment in the N-terminal region of the protein.

**Figure 4 –Figure Supplement 1:**
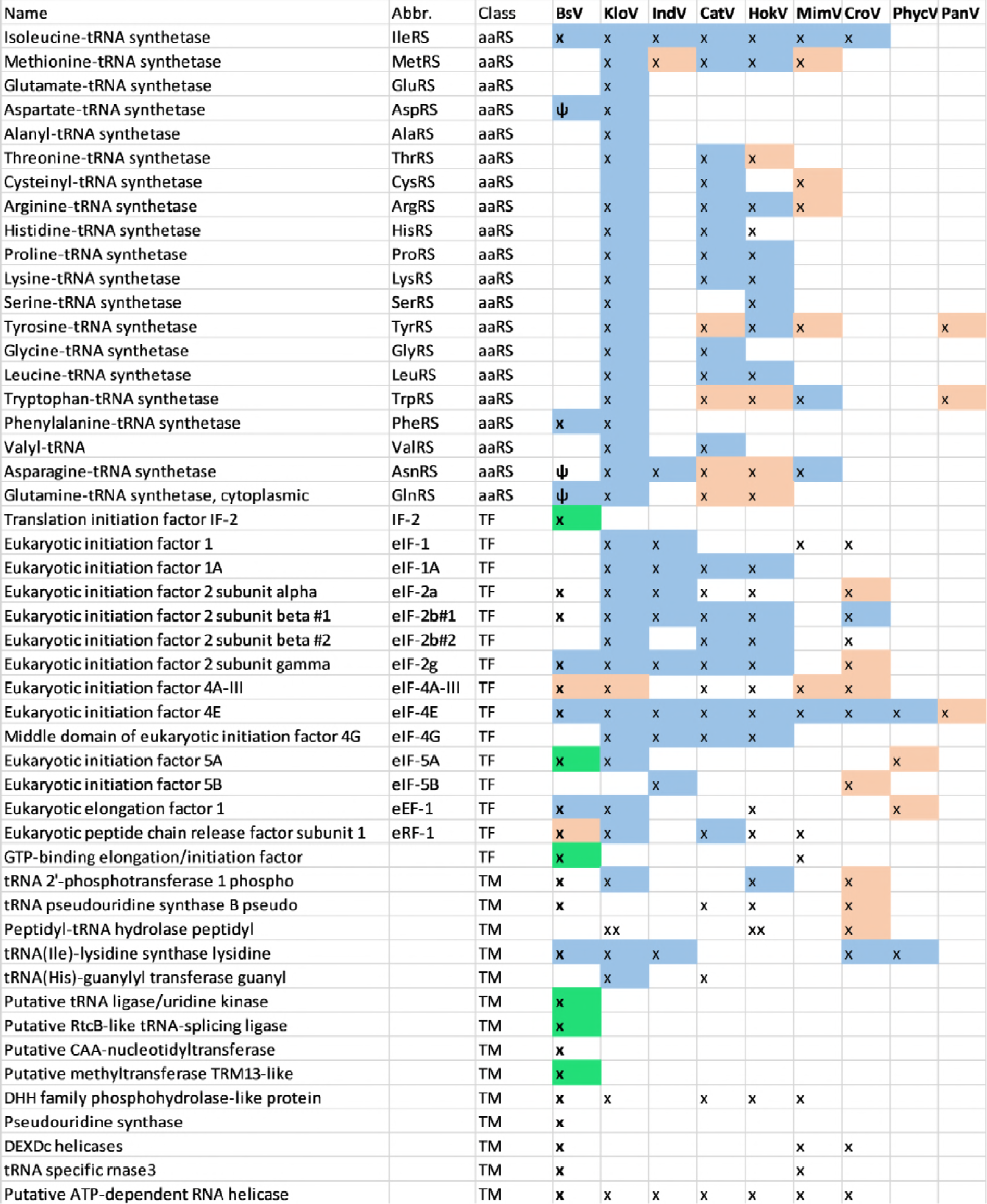
Comparison of translational machinery encoded by giant viruses based on phylogenetic analysis: BsV Translational machinery compared to other NCLDV: blue: monophyletic group, red: recently host acquired, green: recently acquired from bodonid host, ψ: pseudogene

**Figure 5 –Figure Supplement 1:**
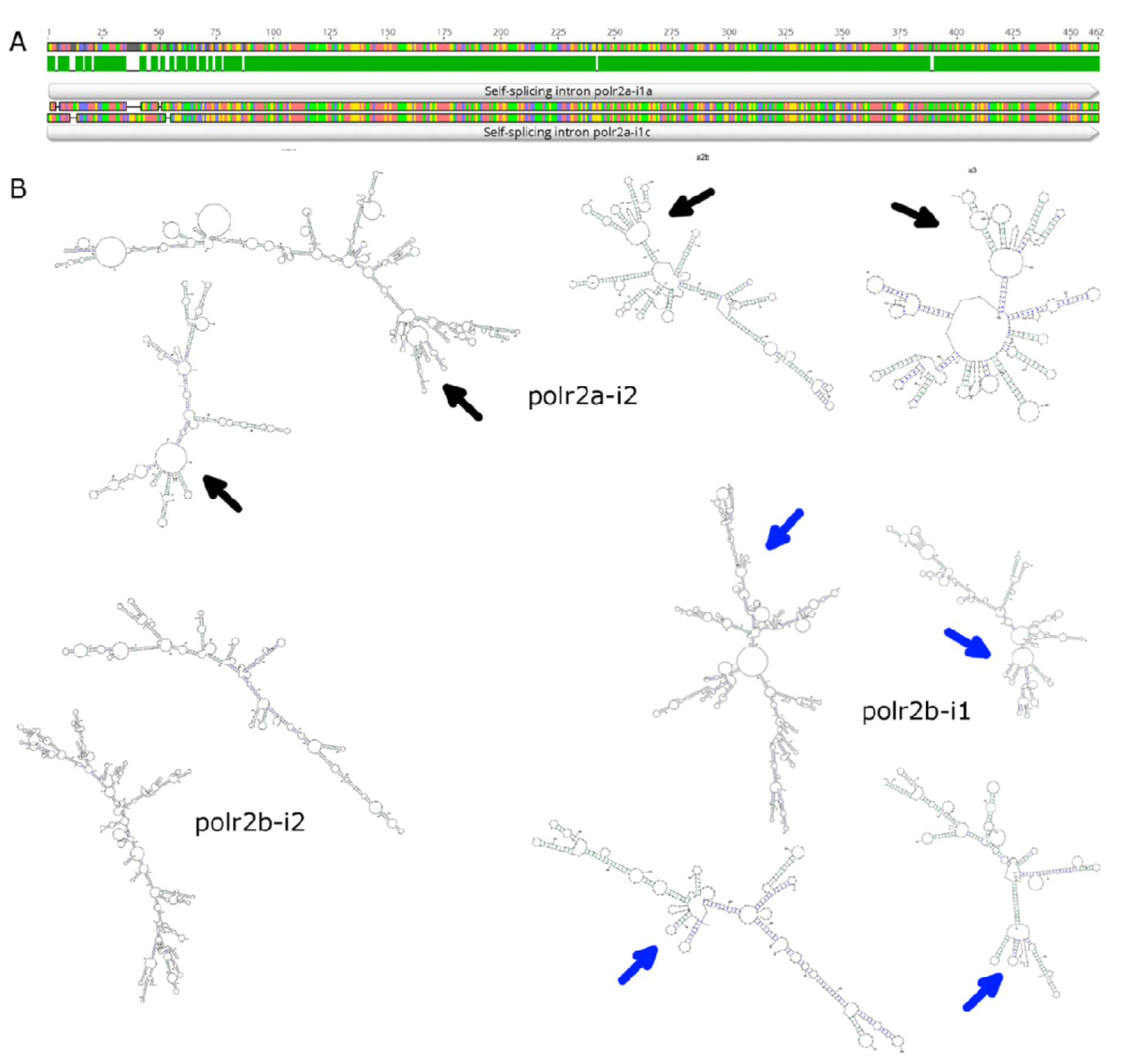
Self-splicing group 1 introns found in BsV. A: DNA Sequence alignment of recently duplicated self-splicing group 1 introns polr2a-i1a and polr2a-i1c (94% DNA sequence ID). B: Secondary structure and conserved putative autocatalytic ribozyme centers of other polr encoded introns.

